# COVID-related anthropause highlights the impact of marine traffic on breeding little penguins

**DOI:** 10.1101/2023.06.30.547199

**Authors:** Benjamin Dupuis, Akiko Kato, Nicolas Joly, Claire Saraux, Yan Ropert-Coudert, Andre Chiaradia, Marianna Chimienti

**Affiliations:** Centre d’Etudes Biologiques de Chizé, UMR 7372 CNRS—La Rochelle Université, 79360 Villiers en Bois, France; Université de Strasbourg, CNRS, Institut Pluridisciplinaire Hubert Curien (IPHC), UMR 7178, 23 rue Becquerel, 67000 Strasbourg, France; Conservation Department Phillip Island Nature Parks, Cowes, VIC 3922, Australia; School of Biological Science, Monash University, Clayton, Australia

**Keywords:** COVID-19 lock-down, Long-term monitoring, Anthropogenic activities, Breeding ecology, Little penguins

## Abstract

The COVID-19 pandemic and its lock-down measures have resulted in periods of reduced human activity, known as anthropause. While this period was expected to be favorable for the marine ecosystem, due to a probable reduction of pollution, shipping traffic, industrial activity and fishing pressure, negative counterparts such as the increased use of disposable plastic and reduced fisheries surveillance and enforcement could counterbalance these positive effects. Simultaneously, on-land pressure due to human disturbance and tourism should have drastically decreased, potentially benefiting land-based marine breeders such as seabirds. Thus, long-term datasets became crucial to differentiate between historical trends and any evident changes resulting from the anthropause. We analyzed 11 years of data on several biological parameters of little penguins (*Eudyptula minor*) from the Penguin Parade ®, a popular tourist attraction at Phillip Island, Australia. We investigated the impact of anthropogenic activities on penguin behavior during the breeding season measured by (1) distribution at sea, (2) colony attendance, (3) isotopic niche (4) chick meal mass, and (5) offspring investment against shipping traffic and number of tourists. The 2020 lock-downs resulted in a near absence of tourists visiting the Penguin Parade ®, which was otherwise visited by 800,000+ visitors on average per year. However, our long-term analysis showed no effect of the presence of visitors on little penguins’ activities. Surprisingly, the anthropause did not triggered any changes in maritime traffic intensity and distribution in the region. While we found significant inter- and intra-annual variations for most parameters, we detected a negative effect of marine traffic on the foraging efficiency. Our results suggest that environmental variations have a greater influence on the breeding behavior of little penguins compared to short-term anthropause events. Our long-term dataset was key to test whether changes in anthropogenic activities affected the wildlife during the COVID-19 pandemic.

**Impact statement:** We found that marine traffic, but not tourist presence, negatively impact the foraging and provisioning behavior of little penguins.

## Introduction

With the development of human activities, ecosystems can no longer be considered as undisturbed and independent entities (Mace, 2014), leading to the concept of socio-ecological systems (Everard, 2020; Wei et al., 2018). Because of the numerous interactions at stake, socio-ecological ecosystems are often complex to analyze (Sugihara et al., 2012). The quasi-continuous presence of humans in most, if not all, ecosystems makes it challenging to understand the full impact of anthropogenic activities on the environment.

In 2020, the COVID-19 pandemic led to periods of lock-downs that resulted in a major reduction of human activities and movement at both local and global level, a period coined as the “anthropause” (Lamers & Student, 2021; Rutz et al., 2020). The anthropause created an opportunity to quantify the impact of human activities on wildlife. To date, studies found both negative and positive effects of this anthropause on wildlife, through for example, increase of predators presence and disturbance on an iconic seabird colony in the Baltic Sea (Hentati-Sundberg et al., 2021), as well as increased species richness in less-disturbed areas (Manenti et al., 2020). Lock-downs also led to increased illegal hunting and plastic pollution, and reduced conservation efforts with negative effects on wildlife (Bates et al., 2021; Kadykalo et al., 2022). In a comparative study, Bates et al. (Bates et al., 2021) showed that despite the decrease in humans’ movement, or industrial activities, the median responses of wildlife to anthropause were centered on 0, because firstly positive and negative effects balanced themselves, while for numerous species, no changes were observed.

Moreover, it can be misleading to consider that the anthropause is a phenomenon homogeneously distributed across the globe. The decrease in human activities was not equal across the planet (Bates et al., 2021). Keeping in mind the level of variation of anthropauses, and eventual anthropulses (i.e. increase of human activities) it is key to study effects of these periods on ecosystems.

It can be complex to study the dynamics of entire ecosystems, specifically within the context of the COVID lock-down, considering the difficulties to carry on with species and habitat monitoring activities during these periods. Monitoring “sentinel species” helps tackling this issue. Sentinel species integrate changes happening across ecosystems’ levels (Durant JM et al., 2009), integrate broader processes into rapidly interpretable metrics, are simpler to study, can respond rapidly to environmental changes and cover a large spatial scale (Bost et al., 2008; Durant JM et al., 2009; Hazen et al., 2019; Siddig et al., 2016). Therefore, long-term dataset on marine predators, especially seabirds, are often used as indicators of ecosystems’ changes (Cairns, 1988; Furness & Camphuysen, 1997; Piatt, Sydeman, et al., 2007).

Data collection via continuous monitoring programs allows researchers to compare the pace of parameters responses to global changes and assess effects of human pressure on wild populations (Cairns, 1988; Durant JM et al., 2009; Einoder, 2009; Ramírez et al., 2017; Tucker et al., 2018, 2019). Techniques used vary depending on the research question and feasibility, comprising of visual observations, counts, nest monitoring, blood sampling and use of bio-logging techniques. In seabirds, chick growth, colony attendance, and individuals’ activity budgets vary at different temporal scales and in relation to both environmental and human activities (Cairns, 1988). Depending on the specific effects of the COVID lock-downs and the relative short period these were put in place, some of these traits might show no responses to anthropogenic activities (Cairns, 1988; Piatt, Harding, et al., 2007).

During breeding season, seabirds are central place foragers exploiting food resources around their breeding colony to which they return due to reproductive requirements (e.g. egg incubation, chick provisioning), hence alternating between nest attendance and foraging trips (Einoder, 2009; Piatt, Sydeman, et al., 2007; Saraux et al., 2011). Seabirds must cope with constraints of living in two different environments, feeding at sea and breeding on land, making them exposed and vulnerable to threats from both land and sea. The little penguin (*Eudyptula minor*) is the smallest penguin species endemic of Australia and New Zealand (BirdLife International, 2023). Phillip Island, Australia, holds one of the largest little penguin colonies in the world with a population estimated between 28,000 and 32,000 individuals (Sutherland & Dann, 2014). The colony located at the “Penguin Parade ®” receives the visit of hundreds of thousands of tourists per year, especially when little penguins return ashore at night (Dann & Chambers, 2013). At sea, little penguins can also interact with maritime traffic such as commercial shipping, recreational or commercial fishing vessels (Cannell et al., 2020; Crawford et al., 2017). Land introduced predators and starvation are the major causes of little penguins’ mortality, but collision with vessels were also reported (Cannell et al., 2016, 2020), even though their foraging range is small (around 30 km for single day trips but can be up to 214 km for multi days trips) (Collins et al., 1999; Poupart et al., 2017; Sánchez et al., 2018).

On land, tourism has been shown to affect various parameters of penguins’ ecology such as stress level, reproductive output (Ellenberg et al., 2007) or behavior (Colombelli-Négrel & Katsis, 2021; Ellenberg et al., 2007; French et al., 2019). At-sea, vessels can directly (Pichegru et al., 2022) or indirectly (Mattern et al., 2013) affect penguins foraging through noise pollution and deterioration of the environment, respectively. During the COVID-19 pandemic, Australia underwent a series of rigid lock-downs, drastically reducing anthropogenic activities. During most of that period, the “Penguin Parade ®” remained closed to the tourists, providing a good opportunity to understand if the anthropause affected ecology of little penguins.

We investigated whether the anthropause affected metrics linked to little penguin’s behavior during the breeding season in 2020 (year with lock-downs) by comparing against 10 years of population monitoring and movement data (2010-2019) to 2020. The studied colony has been monitored for the past 23 years using an automated penguin monitoring system (date, time and weight of penguins recorded when leaving and arriving to the colony), with daily count of penguins arrival at dusk, and with the use of bio-logging techniques (the latter since 2010) (Chiaradia & Kerry, 1999; Ramírez et al., 2015). We tested whether reduced anthropogenic activities influenced little penguins (1) at-sea activity by studying their at-sea distribution, overlap with marine traffic, isotopic diet (in terms of prey type and quantity), and (2) on-land activity by studying their colony attendance (departure and arrival time), and meal size given to their chicks. We considered the daily number of tourists at the Penguin Parade ® as a proxy of land disturbance, and the number of vessels and their overlap with little penguins’ foraging area at-sea as a proxy of the at-sea disturbance. We hypothesized that when land disturbance is reduced during the anthropause, due to the absence of tourists in the parks, little penguins would change their colony attendance pattern by coming and leaving more synchronously, as they will not have to avoid tourist disturbance (Klomp & Wooller, 1991; Rodríguez et al., 2016). Moreover, if the anthropause reduced the at-sea disturbance, little penguins would display a higher foraging efficiency as the overall marine environment and its species will less be disturbed by the traffic, through reduction in noise pollution for instance (Pichegru et al., 2010, 2017).

## Material and Methods

### Study site and long-term monitoring of foraging behavior

The study was conducted on the little penguin breeding colony at Phillip Island, Australia (38°21’S, 145°09’E) from 2010 to 2020. The breeding season of little penguins occurs in the austral spring and summer, from September to December.

For the period of our study (11 breeding seasons, 2010-2020), penguins were captured from their nest boxes and equipped with GPS loggers (Axy-Trek, Italy, Mr Lee, China) recording positions at 120 s interval for incubation and postguard trips and every 20 s for guard trips (table 1). Loggers were attached to their lower backs with Tesa ® tape (Wilson et al., 1997). After returning from their foraging trips, penguins were recaptured at the colony and the logger retrieved. Handling time was kept at less than 5 min. Details of the logger deployment are described in Pelletier et al. (2014), Sanchez et al. (2018) and Barreau et al. (2021) (Barreau et al., 2021; Pelletier et al., 2014; Sánchez et al., 2018). We combined the information obtained from GPS data for estimating the distribution of little penguins at sea, stable isotopes data to investigate their diet, as well as from the automated monitoring system to track changes in body weight and colony attendance.

**Table 1.**
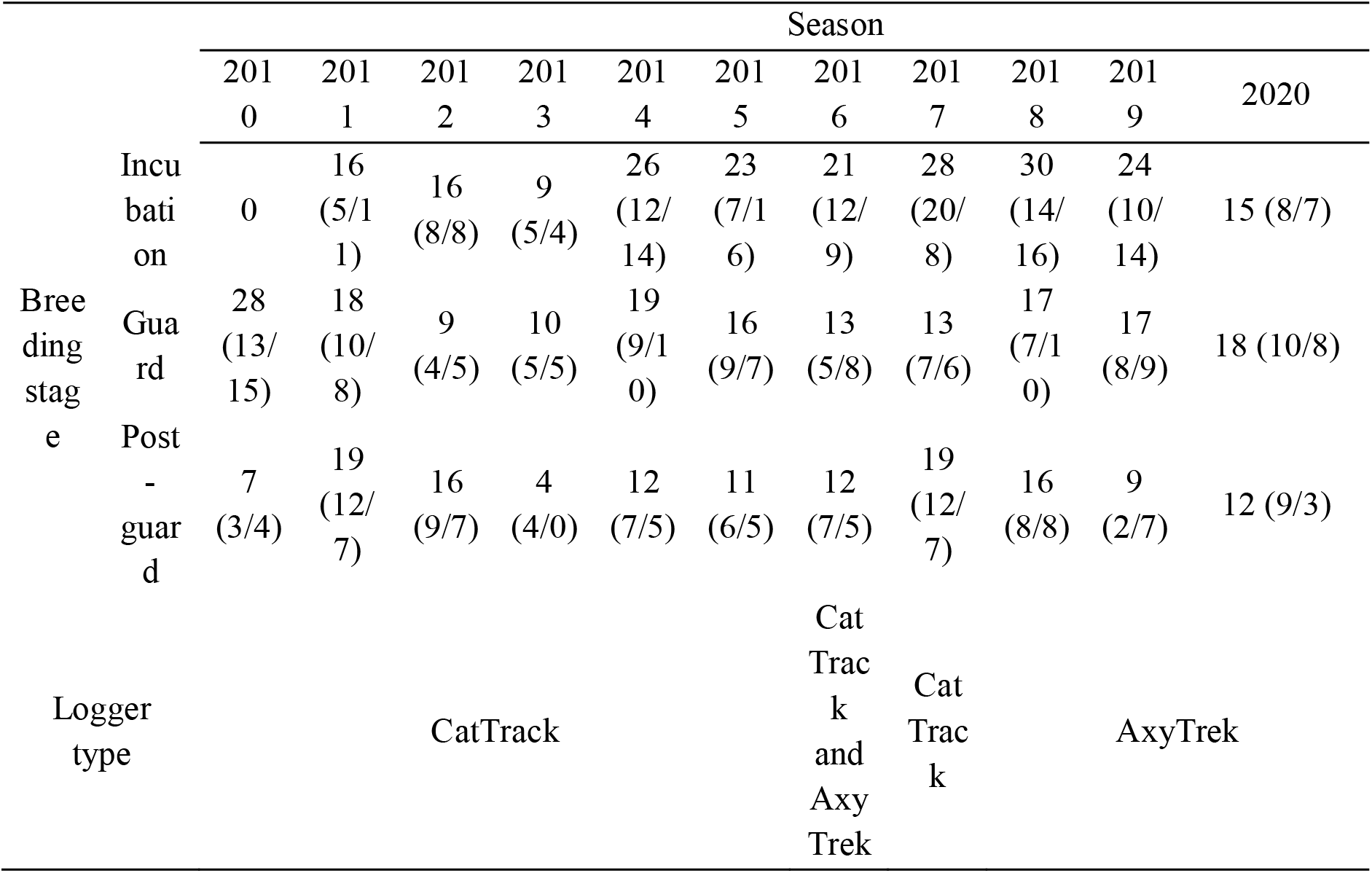
Number of little penguins and type of loggers deployed for each year and breeding stage. (Females/Males)

Two automated penguin monitoring systems (APMS) are placed on the main pathways between the little penguins’ colony and the beach. When walking through APMS, little penguins are individually identified with passive transponders (Allflex, Australia) that had previously been inserted in the back of the penguins, either as chicks or the first time they were encountered in the colony. In addition, APMS record date, time, direction of passage, and the body mass of the individuals (Joly et al., 2022).

This research was conducted under the Phillip Island Nature Parks Animal Experimentation Ethics Committee approval and a research permit issued by the Department of Environment Land, Water and Planning of the state of Victoria, Australia.

### Data manipulation and analysis

Data manipulation and analysis was done in R v4.2.3 (R Core Team, 2023). All scripts used in this analysis are available upon request (Github link removed for anonymity). Unless specified otherwise, results indicate mean and standard error. As well, when more than one variable was considered within a model, all model combinations were tested, and we performed model selection using Akaike’s information criterion (AIC) (Bozdogan, 1987). The model with the lowest AIC was considered as best. Normality of residuals, residual autocorrelation and homoscedasticity were checked graphically. We considered p-values under 0.05 as significant. Unless stated otherwise, pairwise post-hoc comparisons were performed using Holm p-value correction (Holm, 1978).

### GPS data processing

A foraging trip was defined as a period from the departure to the return to the colony. Because little penguins are visual hunters, foraging activities only occur during daylight (Cannell & Cullen, 1998; Chiaradia et al., 2007). Their foraging trips last typically between 1-9 days during incubation (Kato et al., 2008), 1 day during guard (Pelletier et al., 2014) and between 1-17 days during post-guard (Saraux et al., 2011). From each foraging trip, and day out at sea, we extracted a “foraging segment”, intended as the period between nautical dawn and nautical dusk. Therefore, one-day trip contained only one foraging segment, while multiple-days trip could contain several segments. We removed foraging segments with less than 3 GPS locations, and segments starting after sunrise or stopping before sunset, from the analysis. Overall, out of 233 foraging trips, a total of 371 foraging segments were extracted and analyzed (range 1-7 segments per individual).

We calculated the distance between each consecutive location on the WGS ellipsoid using the *pointdistance()* function from the “raster” R package (Hijmans, 2022). Swimming speed was then calculated between two consecutive locations as the distance divided by the time interval. Furthermore, we excluded GPS locations with swimming speed higher than 8 km.h^-1^ (i.e. max swimming speed of little penguins, (Watanuki et al., 2006)), or with a time interval between 2 consecutive GPS locations lower than 7.2 sec (i.e. duplicated points). GPS locations can be obtained only when penguins resurface, therefore it is necessary to interpolate raw GPS data and reconstruct their path. For each foraging segment, we regularized the time interval between each location by performing spatial interpolation at 15-min interval using the correlated random walk algorithm within the *crawlWrap()* function from “MomentuHMM” R package (McClintock & Michelot, 2018).

Interpolated foraging segments were projected into the GDA94 / Australian Albers projection. For each breeding season, we then used the *kernelUD()* function from the “adehabitatHR” R package (Calenge, 2006) to calculate 50% (core area), and 95% (homerange) kernel utilization distribution (UD). The smoothing parameter *h* was calculated using the ad hoc method (Seaman et al., 1998).

### Stable isotope data processing

To describe the isotopic niches of little penguins and examine differences between 2020 and previous years (2010-2019), we analyzed δ15N and δ13C stable isotopes from 842 blood samples (n = 196 in incubation, n = 367 in guard, n = 279 in post-guard). Values in δ15N increase with prey trophic level, while δ13C values are higher inshore than offshore (Hobson et al., 1994). We followed the protocol described in Chiaradia et al. (Chiaradia et al., 2016). Whole blood was freeze-dried and then powdered. As mass C/N ratios were all below 3.5, there was no need for correction of lipid contents in whole blood (Post et al., 2007). Isotopic analysis was then performed by means of a Robo-Prep elemental analyzer coupled to a Europe 20:20 continuous-flow isotope ratio mass spectrometer. Based on replicate measurements of within-run standards, measurement error was estimated to be ±0.3 and ±0.1‰ for δ15N and δ13C measurements, respectively.

### Automated penguin monitoring system

We evaluated two measures of body mass variation. First, we calculated the mass gained after a foraging trip, which we considered to be an estimate of foraging efficiency (Saraux et al., 2011). Two body masses were considered belonging to the same foraging trip when their records were consecutive in date and time for a given transponder number and the trip duration was not longer than 1 d in guard and 17 d in incubation and post-guard (Salton et al., 2015). Then, for post-guard only, we calculated the overnight mass variation after returning from a foraging trip, which we considered to be an estimate of chick provisioning during chick-rearing. During this stage, little penguins stay only a few hours in the colony, so we assumed that all body mass loss was due to chick provisioning. Body mass gained at sea was only considered when ranging from 700 to 1700 g and body mass change from -75 to 500 during incubation and 0 to 600 g during guard and post-guard (Joly et al., 2022; Saraux et al., 2011).

Using APMS, we also calculated penguins’ attendance to the colony. When penguins crossed the weighbridge, it registers the timestamp, and transponder number of the penguin, allowing us to now departure and arrival times of each foraging trips. We calculated departure and arrival times relative to nautical dawn and dusk, respectively, to account for variation in day length (Rodríguez et al., 2016).

### Proxies of anthropogenic activities

Given that little penguins breed on land and forage at sea, we defined both on land and at sea indicators of anthropogenic activities. The number of tourists present each night was used as an index of human activity on land. This number was monitored daily between 2010 and 2020. Over the studied period, artificial lighting (orange halogen lights, 3 lux) was used to enhance visibility of penguins for tourists. These lights were turned on from sunset to 1.5 h after the arrival of the first penguins (Rodríguez et al., 2016). During the COVID lock-downs, these lights were still in place but without the presence of tourists.

For the activity at sea, we used the number of vessels (fishing, commercial and leisure) within the little penguin foraging area (longitude 145 to 146°E, latitude 38.5 to 39.5°S) during their breeding season (September to December). We used the open-source dataset from the Australian Marine Safety Authority (https://www.operations.amsa.gov.au/Spatial/DataServices/DigitalData) and for each vessel we obtained its ID, latitude, longitude, type and timestamp. As vessels transmit their locations at different time interval (from one per 15 minutes to once a day), we built daily indices by keeping only one location per vessel and day: the closest to noon available. Data were available only between 2014 and 2020. Information earlier than 2014 was not considered because of the lower time resolution compared to later data, and data for November 2019 was missing. Hereafter, we refer to the number of vessels within little penguin foraging grounds as the marine traffic intensity. We calculated marine traffic UD using the same method described before for the little penguins.

### Statistical analysis

#### Variation of anthropogenic activities

Using linear models, we investigated the variation of the number of vessels in little penguins foraging area and the number of tourists at the penguin parade ® between months and years. Then, using pairwise post-hoc comparison, we tested the difference between the COVID year (2020) and the previous years.

#### Spatial variation of at sea distribution and overlap with marine traffic

Overlap analyses were performed using the Utilization Distribution Overlap Index (UDOI, (Fieberg & Kochanny, 2005)) which quantifies the pattern of space-use as a function of the product of the overlapping UDs. UDOI is equal to 0 when two UDs do not overlap and to 1 if the UDs are completely overlapping and uniformly distributed. Values higher than 1 indicate higher normal overlap relative to uniform space-use. UDOI were calculated using the *kerneloverlap()* function from the “adehabitatHR” R package (Calenge, 2006).

We calculated the UDOIs of the 95% UD of the at sea distributions of penguins for all years. We calculated the UDOIs of a year A with all the other years, generating a distribution of UDOIs for year A. We then assessed whether the observed distribution in 2020 was different compared to the previous years. We also calculated the UDOI between little penguins and marine traffic. Again, we calculated UDOIs of year A (little penguin) with all the available years (marine traffic). We obtained distributions of the UDOIs between little penguins and marine traffic. We tested differences between years using generalized linear models (GLMs) with a gamma distribution. We then performed post-hoc pairwise comparison to assess the significance of differences observed between 2020 and the other years.

#### Effect of number of tourists on little penguins attendance and foraging efficiency

Linear models (LMs) were used to test the effect of lock-down on (a) average departure and arrival times of little penguin relative to nautical dawn and nautical dusk, respectively and (b) average mass variation per day over a foraging trip, and overnight. Models were built using the ‘nlme’ R package (Pinheiro et al., 2021). For both models (a and b) we considered number of tourists and number of vessels as explanatory variables, and breeding stage and season as fixed effects. To asses the effect sizes of both vessels and tourists counts, we standardized these data (see equation 1).

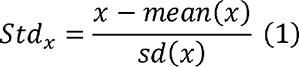

#### Quantification of isotopic niche

We computed standard ellipse area corrected for small sample size and extreme values (SEA_C_) to estimate isotopic niche width and overlap among the different years and breeding stages. SEA_C_ represents the isotopic niche width of 40% of typical individuals within the groups, based on bivariate normal distribution. The overlap in SEA_C_ was calculated for all pairs of years within a breeding stage following (Catry et al., 2016) where isotopic niche overlap was expressed as a proportion of the area of overlap between two SEA_C_ to its own SEA_C_. We also computed Bayesian Standard ellipse area (SEA_B_) (n = 20000 iterations) to obtain credible intervals (99%, 95% and 50%) for the calculated ellipses. We considered non-overlapping 95% CI around SEA_B_ as an indicator of statistically significant difference between niches width. For all this analysis, we followed the method described in (Jackson et al., 2011) and the ‘SIBER’ R package.

## Results

### Variation of anthropogenic activities

In 2020, during the COVID lock-downs, Phillip island nature park remained closed for most of the breeding season, resulting in number of tourists 10 times lower than usual (180.4 ± 27.9 tourists per day in 2020 vs. 1770.0 ± 20.5 on average in 2010-2019, all *p* < 0.001, figure 1, supplementary table 1.A and 2.A).

**Figure 1:**
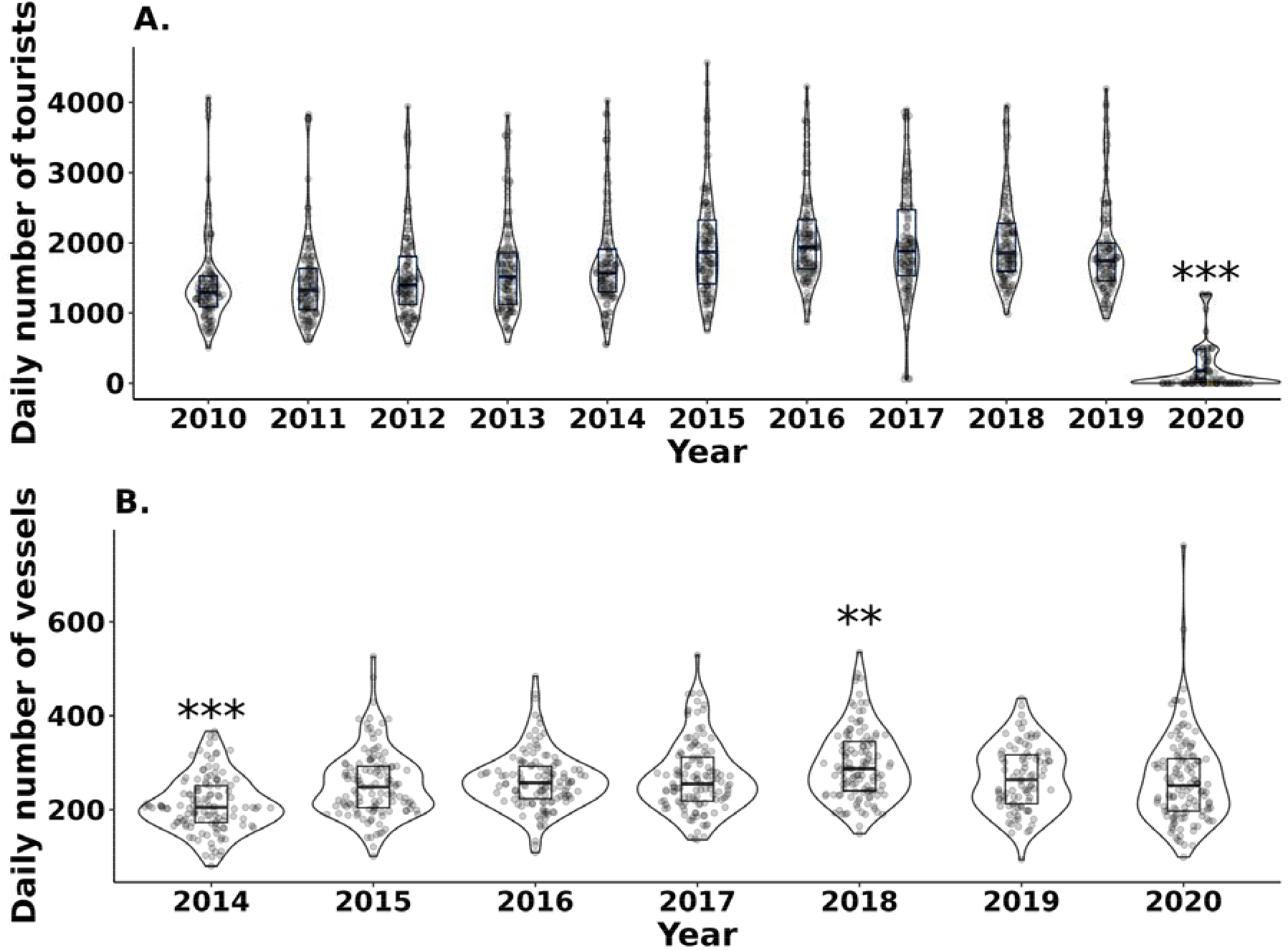
Evolution of anthropogenic activities in the studied area. (A) Daily number of tourists at the penguin parade ® between 2010 and 2020. (B) Daily number of vessels at sea in the foraging area of little penguins between 2014 and 2020. Astariscs represent statistical significance of post-hoc comparisons between 2020 (lock-down season) and the others (2010-2019, ** = *p* < 0.01, *** = *p* < 0.001).

In 2020, the daily average number of vessels recorded at sea was 262 ± 8.51. This was significantly higher than the one recorded for 2014 of 212 ± 5.35 (estimate = 50.607, t = 5.562, *p* < 0.001), and lower than 2018 with 297 ± 6.91 vessels (estimate = -34.279, t = -3.767, *p* = 0.002, figure 1, supplementary table 1.B and 2.B).

### Spatial variation of at sea distribution and overlap with marine traffic

While spatial distribution of marine traffic remained similar across seasons, little penguins core (50% UD) and home ranges (95% UD) showed great inter-annual variation across the studied period (figure 2). We compared the overlaps of penguins distribution in 2020 (average UDOI of 0.86 ± 0.07) to all the other years (average UDOI ranging from 0.58 ± 0.09 in 2015 in to 1.09 ± 0.07 in 2014) at 95% UD. We did not find any significant difference in the overlap distributions in 2020 vs any other season (supplementary table 1.C and 2.C).

**Figure 2:**
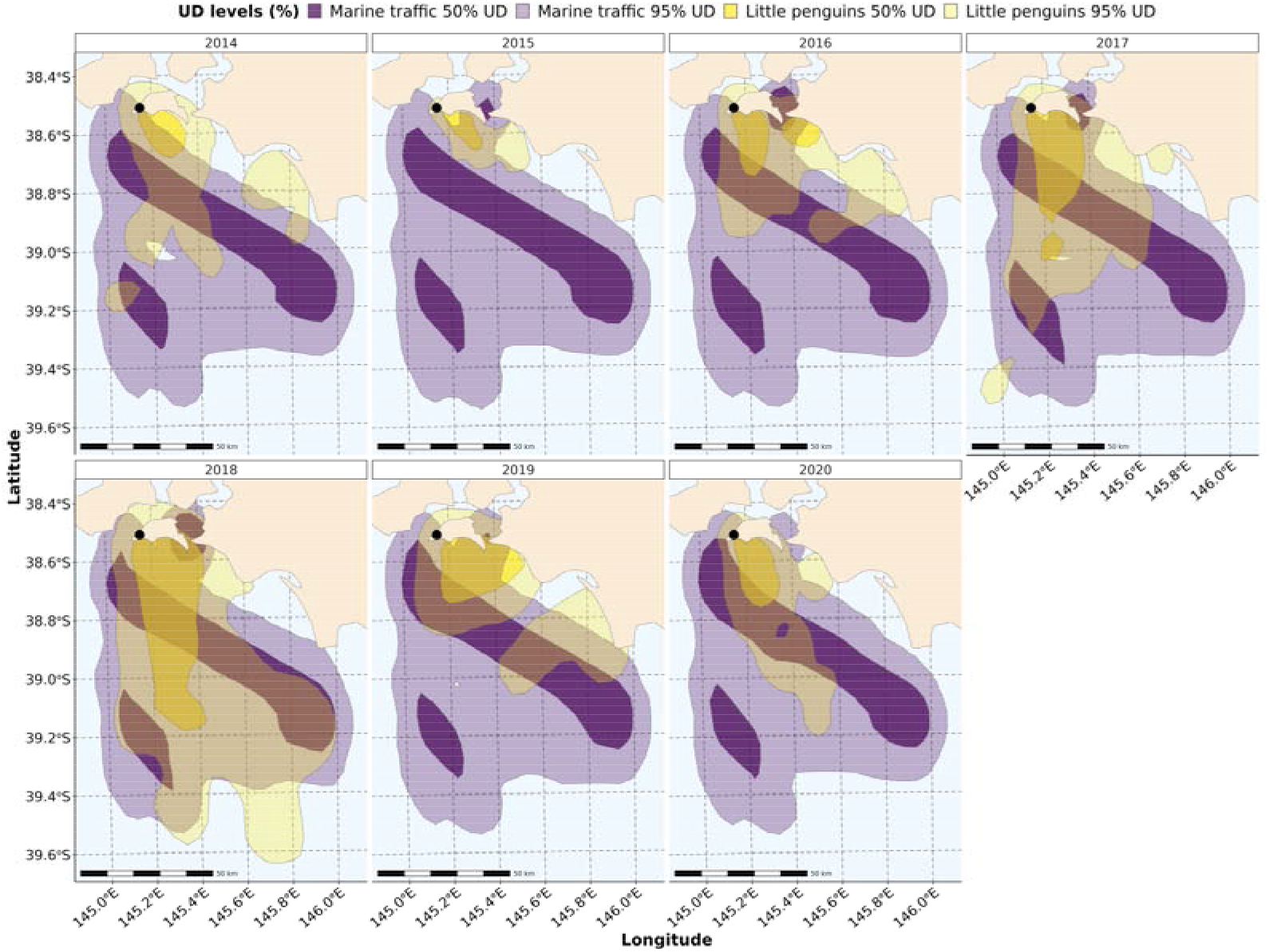
Spatial distribution of little penguins and marine traffic between 2014 and 2020. The black dot represents the studied colony in Phillip Island.

Model selection pointed at the model with the effect of the *year* as explanatory variable as best (supplementary table 1.D). We found variation in the overlap between marine traffic and little penguins’ distributions (from 2014 to 2020). In 2020, the overlap between little penguins and marine traffic was significantly lower (0.184 ± 0.009) than in 2018 (average = 0.649 ± 0.144, estimate = 3.879, *p* < 0.001) and 2017 (average = 0.358 ± 0.128, estimate = 2.627, *p* < 0.001), but higher than the one of 2015 (average = 0.021 ± 0.003, estimate = - 41.868, *p* <0.001, supplementary table 2.D).

### Effect of number of tourists and marine traffic on colony attendance

Little penguins left the colony on average 52.9 ± 0.5 minutes (n = 11116) before nautical dawn and there is no difference across seasons. The best model testing the effect of anthropogenic activities on the time of departure relative to nautical dawn retained only the effect of breeding stage as explanatory variable (supplementary table 1.E, p-value <0.01), with therefore no significant inter-annual variations (supplementary table 2.E). During incubation, penguins left 31.8 minutes (95% CI [24.5; 39.0]) before nautical dawn, compared to 74.9 minutes (95% CI [67.7;82.2]) during guard and 47.1 minutes (95% CI [39.9;54.4]) during post-guard (figure 3A).

**Figure 3:**
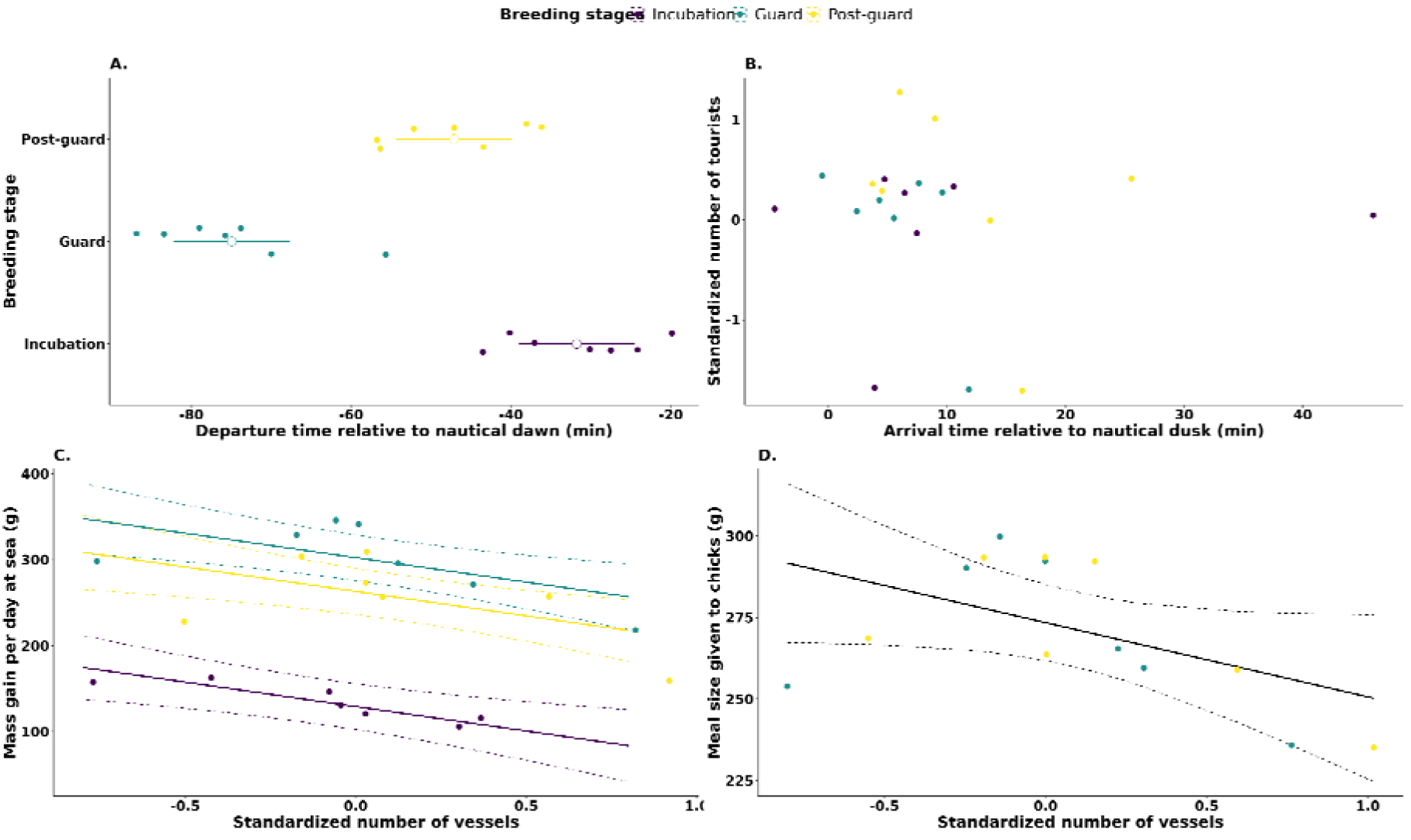
Anthropogenic activities effect on APMS-derived parameters. (A) Departure time relatively to nautical dawn, and (B) Arrival time relatively to nautical dusk of little penguins at the colony depending on the number of tourists. Effect of the number of vessels on the (C) Mass gain per day at sea and (D) Meal mass given to the chicks at the colony. Colored points represent a season average, and white points the overall mean with its SE. Dashed lines represent the 95% CI around model predictions.

Model selection for the models testing the effect of anthropogenic activities on the time of arrival relative to nautical dusk pointed at the null model as best (supplementary table 1.F), indicating an absence of effect of tourists presence and marine traffic on colony attendance. Penguins showed highly synchronized arrival time regardless of season or breeding stage, arriving at the colony on average 8.2 ± 0.4 min after nautical dusk (n = 11087, figure 3B, supplementary table 2.F).

### Effect of number of tourists and marine traffic on foraging efficiency

Over a foraging trip, penguins gained on average 258.65 ± 1.75 g per day (n = 6617). The best model testing the effect of anthropogenic activities and temporal variations on mass gained per day at sea retained breeding stage and daily average number of vessels at sea as explanatory variables (figure 3, supplementary table 1.G), indicating an effect of marine traffic intensity but not of anthropause on foraging efficiency. Higher number of vessels was associated with lower mass gain at sea for little penguins (estimate = - 56.9 ± 17.3 g, *F* = 10.8, *p* = 0.004, supplementary table 2.G). Predicted breeding stage mass gain were all significantly different from one another (*F* = 46.6, all *p* < 0.05). During incubation, penguins gained 127.4 g (95% CI [100.6;154.3]) per day, compared to 300.7 g (95% CI [274.2;327.1]) in guard, and 261.4 g (95% CI [234.6;288.2]) in post-guard.

The average overnight mass change during post-guard, i.e. meal size, was of 278.6 ± 0.1 g (n = 1794). Though the best model was the one with the average number of vessels at sea (supplementary table 1.H), its effect was not significant on meal size given to the chicks (estimate = - 22.8 ± 11.2, *F* = 4.2, *p* = 0.06, figure 3D, supplementary table 2.H).

### Quantification of isotopic niche

A total of 842 blood samples were collected from little penguins across the different breeding stages (supplementary table 3). We observed variations in the isotopic niche values and areas at different breeding stages over 10 years (figure 4). During incubation, the SEAb mode of 2020 was 0.79 ‰² with 95% CI [0.48;1.25] and was significantly higher than 2 other years, 2011 (0.19 ‰² [0.13;0.31]) and 2015 (0.21 ‰² [0.14;0.34] (figure 5). During the guard stage, SEAb was higher for 2020 (0.98 ‰² [0.69; 1.39] than 2011 again (0.24 ‰² [0.17;0.33]), and 2010 (0.46 ‰² [0.33;0.65]. Finally, during the post-guard, the SEAb of 2020 decreased (0.60 ‰² [0.45;0.87]). It was still significantly higher than the SEAb of 2011 (0.29 ‰² [0.21;0.42]), but also significantly lower than the one of 2014 (1.51 ‰² [0.88; 2.72]). These inter annual variations lead to low overlap between the isotopic niches of little penguins between 2010 and 2020 (table 2).

**Figure 4:**
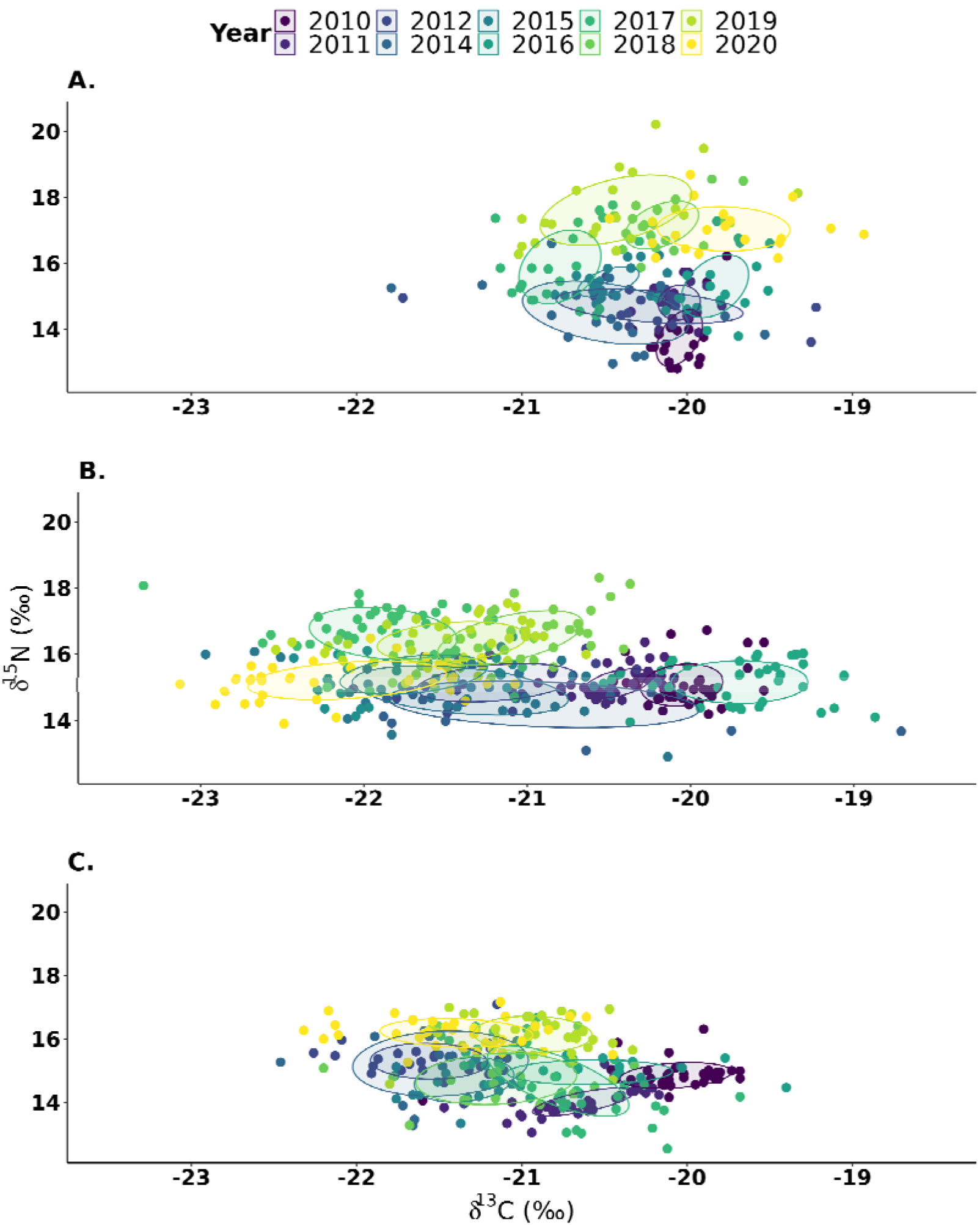
Isotopic niche of little penguins between 2010 and 2020, across different breeding stages. Ellipses represent the corrected standard ellipses of each niche (40% of the individuals).

**Figure 5:**
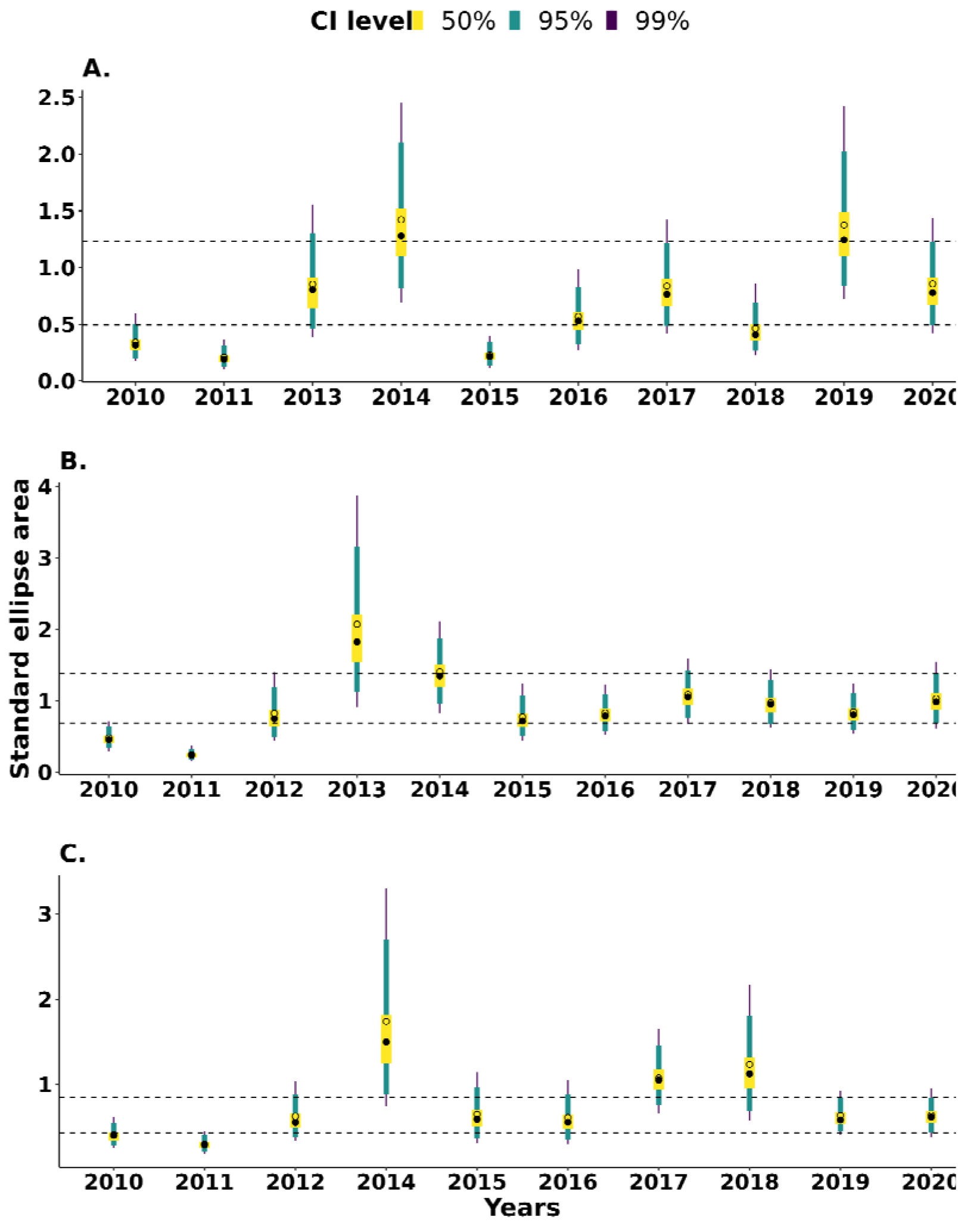
Standard ellipses’ area of little penguin’s isotopic niches between 2010 and 2020 during different breeding stages. Black dots represent the mode of the Bayesian standard ellipse area, and error bars the confidence intervals at 50, 95 and 99%. Circles represent the corrected standard ellipse areas.

## Discussion

Humans have increasingly altered natural habitats, triggering changes in movements, habitat use and population dynamics in wild species (Duhem et al., 2008; Holles et al., 2013; Margalida et al., 2014). The anthropause period caused by the COVID-19 pandemic set an unprecedented opportunity to study the effects of reduced human activities on the biology and ecology of a range of species (Rutz et al., 2020). During the anthropause, human activity on land decreased massively in our studied area, with a reduction almost by a factor 10 in the number of tourists of the Penguin Parade ®, Phillip Island Nature Park, Australia. Contrary to the expected (Bates et al., 2021), the lock-down policy did not seem to affect the marine traffic, neither spatially nor quantitatively within the penguin foraging zone within our study site in Bass Strait. This specific setup allowed us to specifically study the effect of on land activity through a stable at-sea potential pressure throughout the study period. Despite the important inter-annual variability in at-sea distribution and diet of little penguins over the studied period (2010-2020), no effect of the anthropause was found. Still, we found anthropogenic effect not linked with the anthropause. Despite the marine traffic intensity stability over the studied period, thanks to our long-term data set, we were able to identify a negative relationship between marine traffic intensity and mass gained at sea per day by little penguins.

Human activities are known to affect seabirds’ physiology and behaviour. Previous studies showed the negative effects of anthropogenic noise (Pichegru et al., 2017), human presence (Ellenberg et al., 2006, 2013), domestic animal (Ratcliffe et al., 2010), food waste (Grémillet et al., 2008) and marine pollution (Trathan et al., 2015) on seabirds. Studies with similar conditions to our study (i.e. seabird in parks or area without tourists due to lock-downs) found that the absence of tourists could, counter-intuitively, lead to more disturbance for the seabirds, which translated in a later laying date and more egg predation (Hentati-Sundberg et al., 2021), underlining the protective role tourists can have for some species. We did not find such an effect in the parameters studied in this paper. Here, we found no effects of the presence of tourists on little penguins activities.

Still, little penguins are known to be sensitive to anthropogenic activities like human presence around the nest (Colombelli-Négrel & Katsis, 2021). A recent study identified the negative effect of anthropogenic activities, *i.e.* white light sources, at night on little penguins (Costello & ColombelliDNégrel, 2023), but evidences are mixed since other study find the opposite (Rodríguez et al., 2018). Multiple hypotheses could explain the absence of response during the anthropause in our study. One could argue that the duration and/or magnitude of the anthropause was negligible to trigger a response in the foraging behavior of little penguins. Plasticity being species dependent (Crawford et al., 2017), more studies on little penguins would be necessary to assess the extent of their plasticity in response to anthropogenic activities, and the potential different threshold that could trigger a response in the studied parameters (Cairns, 1988). On land, predators of little penguins are mainly goannas, snakes and cats (Colombelli-Négrel & Katsis, 2021). However, these predators are a not a thread at Phillip Island, thanks to a successful conservation program in place, with a well managed tourists’ pressure (BirdLife International, 2023; Rodríguez et al., 2018). Long-term exposition to tourists at the Penguin Parade ®, little penguins could have habituated to anthropogenic disturbance (Rodríguez et al., 2016; Viblanc et al., 2012). Therefore it could mean that the predation and disturbance pressure on-land were unchanged during the lock-downs. Finally, the absence of shifts on little penguins trophic niche in 2020 suggests that even if the marine benefited from the lock-down, improving food availability, we did not detected any changes in the diet of little penguins.

To our knowledge, this paper provides the first empirical assessment of the negative effect of marine traffic on little penguins’ foraging behavior during their breeding period. Our study underlines that the at-sea disturbances are more important than the on-land ones when it comes to affect little penguins foraging. The lock-downs did not trigger a reduction of the marine traffic in the Bass Strait but thanks to a long-term data set, we were able to asses the effect of marine traffic on little penguins foraging. As expected, we found a negative effect of marine traffic intensity on little penguins foraging efficiency but not on their spatial distribution.

Our spatial analysis revealed an overlap between little penguins and marine traffic in the Bass strait. However, in the Bass strait, fisheries represent only a small proportion (< 1 %) of the marine traffic in comparison with cargo (50-60 %), tanker (10-20 %), and passenger vessels (5-8 % supplementary figure 1). It is therefore unlikely that the observed effect is due to a competition with fisheries for food. Marine traffic can cause other indirect disturbance, through avoidance behaviour of predators (Jarrett et al., 2021; Pichegru et al., 2022) and preys (Ivanova et al., 2020), or environmental pollution. More investigations is needed on the mechanism to fully understand how marine traffic impacts little penguins to be implemented on marine spatial planning.

Using quantitative information about human activity, like the number of tourists, rather than qualitative one (e. g. comparing lock-down season vs past observed trends) is key to compare study results and assess the shape of the response of wildlife to anthropogenic activities. Indeed, many studies fail to properly quantify anthropogenic activities, and only compared “COVID years” with other years before and/or after (Gordo et al., 2021; Hentati-Sundberg et al., 2021; Sala et al., 2021). While informative, this approach does not allow to properly disentangle anthropogenic pressure from seasonal and environmental variations. Incorporating a quantification of anthropogenic pressure in our models (i.e. number of tourists and vessels) allowed us to disentangle the natural inter- and intra-annual variations from anthropogenic pressure. Our study highlighted that long-term monitoring studies are key to be able to disentangle such effects.

The effect of the anthropauses caused by the COVID related lock-downs on little penguins’ ecology during the breeding might be negligible compared to the ones induced by long-term environmental variations and global changes (Joly et al., 2022). Other significant effects found in our study are mostly related to intra and inter-annual variations. Thanks to long-term monitoring and online data availability, we were able to have a detailed picture of the impact of anthropogenic activities over the 10 years period. Species showing high plasticity and therefore quickly responding to reduced pressures during anthropauses are likely to use that same plasticity in the other way when anthropogenic activities increase again. Such punctual changes could be buffered by phenotypic plasticity and unlikely to change population trends compared to long-term variations (Gordo et al., 2021).

In conclusion, we did not detected any positive or negative effect of COVID-19 lock-downs on the little penguin breeding ecology, despite of our robust dataset used in the analysis. We did show that behavioral variations during their breeding cycle of little penguins of the Phillip Island Nature Park was mostly due to the inter- and intra-annual variation. We revealed that anthropogenic effect due to increased marine traffic can affect foraging efficiency of little penguins. Still, as seabirds live at the interface between sea and land, more information needs to be gathered on the mechanisms behind the effect of marine activity on little penguins foraging and on the effect of on-land anthropogenic activity on little penguins breeding. Better understanding these sources of pressure could help the efficient implementation of marine spatial planning and validate the efficiency of mitigation measures occurring in natural parks. Given the important specific and spatial variability in the responses to anthropogenic activities, the fast change of the marine environment in this region, maintaining and developing long-term monitoring sites and studies is key to guide conservation policies. This will help researchers to better distinguish between environmental and anthropogenic effects on wild species.

## Authors contributions

Conceptualization: AC, YRC, AK

Data curation: BD, MC, NJ, AK

Analysis: BD

Writing: BD, MC

Review of draft: all authors

Supervision: MC, AK, AC

## Acknowledgements

We thank Phillip Island Nature Parks for their continued support and commitment to penguin research. We thank all the students and volunteers for their tireless support in collecting these data over the years, particularly the Nature Parks’s research technical staff Leanne Renwick, Paula Wasiak, Meagan Tucker, Marjolein van Polanen Petel and Jordan Roberts. We thank all the students and volunteers for their tireless support in collecting these data. Field work protocol was approved by the ethics committee of the Phillip Island Nature Park Animal Experimentation Committee with a research permit issued by the Department of Environment, Land, Water and Planning of Victoria, Australia.

## Data availability statement

Sample dataset and all R codes used to manipulate and analyze the data used are available at https://github.com/bendps/COVIDxLP.

## References

Barreau, E., Kato, A., Chiaradia, A., & Ropert-Coudert, Y. (2021). The consequences of chaos: Foraging activity of a marine predator remains impacted several days after the end of a storm. PLOS ONE, 16(7), e0254269. https://doi.org/10.1371/journal.pone.0254269

Bates, A. E., Primack, R. B., Biggar, B. S., Bird, T. J., Clinton, M. E., Command, R. J., Richards, C., Shellard, M., Geraldi, N. R., Vergara, V., Acevedo-Charry, O., Colón-Piñeiro, Z., Ocampo, D., Ocampo-Peñuela, N., Sánchez-Clavijo, L. M., Adamescu, C. M., Cheval, S., Racoviceanu, T., Adams, M. D., … Duarte, C. M. (2021). Global COVID-19 lockdown highlights humans as both threats and custodians of the environment. Biological Conservation, 263, 109175. https://doi.org/10.1016/j.biocon.2021.109175

BirdLife International. (2023). Species factsheet: Eudyptula minor. Downloaded from http://datazone.birdlife.org/species/factsheet/little-penguin-eudyptula-minor on 30/06/2023

Bost, C.-A., Jaeger, A., Huin, W., Koubbi, P., Halsey, L., Hanuise, N., & Handrich, Y. (2008). Monitoring prey availability via data loggers deployed on seabirds: advances and present limitations. Fisheries for Global Welfare and Environment Memorial Book of the 5th World Fisheries Congress, 121–137.

Bozdogan, H. (1987). Model selection and Akaike’s Information Criterion (AIC): The general theory and its analytical extensions. Psychometrika, 52(3), 345–370. https://doi.org/10.1007/BF02294361

Cairns, D. K. (1988). Seabirds as Indicators of Marine Food Supplies. Biological Oceanography, 5(4), 261–271. https://doi.org/10.1080/01965581.1987.10749517

Calenge, C. (2006). The package adehabitat for the R software: tool for the analysis of space and habitat use by animals. Ecological Modelling, 197, 1035.

Cannell, B. L., Campbell, K., Fitzgerald, L., Lewis, J. A., Baran, I. J., & Stephens, N. S. (2016). Anthropogenic trauma is the most prevalent cause of mortality in Little Penguins, Eudyptula minor, in Perth, Western Australia. Emu - Austral Ornithology, 116(1), 52–61. https://doi.org/10.1071/MU15039

Cannell, B. L., & Cullen, J. M. (1998). The foraging behaviour of Little Penguins Eudyptula minor at different light levels. Ibis, 140(3), 467–471. https://doi.org/10.1111/j.1474-919X.1998.tb04608.x

Cannell, B. L., Ropert-Coudert, Y., Radford, B., & Kato, A. (2020). The diving behaviour of little penguins in Western Australia predisposes them to risk of injury by watercraft. Aquatic Conservation: Marine and Freshwater Ecosystems, 30(3), 461–474. https://doi.org/10.1002/aqc.3272

Catry, T., Lourenço, P. M., Lopes, R. J., Carneiro, C., Alves, J. A., Costa, J., Rguibi-Idrissi, H., Bearhop, S., Piersma, T., & Granadeiro, J. P. (2016). Structure and functioning of intertidal food webs along an avian flyway: a comparative approach using stable isotopes. Functional Ecology, 30(3), 468–478. https://doi.org/10.1111/1365-2435.12506

Chiaradia, A., & Kerry, K. R. (1999). Daily Nest Attendance and Breeding Performance in the Little Penguin Eudyptula Minor at Phillip Island, Australia. 8.

Chiaradia, A., McBride, J., Murray, T., & Dann, P. (2007). Effect of fog on the arrival time of little penguins Eudyptula minor: a clue for visual orientation? Journal of Ornithology, 148(2), 229–233. https://doi.org/10.1007/s10336-007-0125-5

Chiaradia, A., Ramírez, F., Forero, M. G., & Hobson, K. A. (2016). Stable Isotopes (δ13C, δ15N) Combined with Conventional Dietary Approaches Reveal Plasticity in Central-Place Foraging Behavior of Little Penguins Eudyptula minor. Frontiers in Ecology and Evolution, 3, 154. https://doi.org/10.3389/fevo.2015.00154

Collins, M., Cullen, J. M., & Dann, P. (1999). Seasonal and annual foraging movements of little penguins from Phillip Island, Victoria. Wildlife Research, 26(6), 705–721. https://doi.org/10.1071/wr98003

Colombelli-Négrel, D., & Katsis, A. C. (2021). Little penguins are more aggressive on islands that experience greater unregulated human disturbance. Animal Behaviour, 182, 195–202. https://doi.org/10.1016/j.anbehav.2021.10.012

Costello, E. C., & ColombelliDNégrel, D. (2023). Human activities at night negatively impact Little Penguin (Eudyptula minor) numbers and behaviours. Ibis, ibi.13217. https://doi.org/10.1111/ibi.13217

Crawford, R., Ellenberg, U., Frere, E., Hagen, C., Baird, K., Brewin, P., Crofts, S., Glass, J., Mattern, T., Pompert, J., Ross, K., Kemper, J., Ludynia, K., Sherley, R. B., Steinfurth, A., Suazo, C. G., Yorio, P., Tamini, L., Mangel, J. C., … Small, C. (2017). Tangled and drowned: a global review of penguin bycatch in fisheries. Endangered Species Research, 34, 373–396. https://doi.org/10.3354/esr00869

Dann, P., & Chambers, L. (2013). Ecological effects of climate change on little penguins Eudyptula minor and the potential economic impact on tourism. Climate Research, 58(1), 67–79. https://doi.org/10.3354/cr01187

Duhem, C., Roche, P., Vidal, E., & Tatoni, T. (2008). Effects of anthropogenic food resources on yellow-legged gull colony size on Mediterranean islands. Population Ecology, 50(1), 91–100. https://doi.org/10.1007/s10144-007-0059-z

Durant JM, Hjermann DØ, Frederiksen M, Charrassin JB, Le Maho Y, Sabarros PS, Crawford RJM, & Stenseth NC. (2009). Pros and cons of using seabirds as ecological indicators. Climate Research, 39(2), 115–129. https://www.int-res.com/abstracts/cr/v39/n2/p115-129/

Einoder, L. D. (2009). A review of the use of seabirds as indicators in fisheries and ecosystem management. Fisheries Research, 95(1), 6–13. https://doi.org/10.1016/j.fishres.2008.09.024

Ellenberg, U., Mattern, T., & Seddon, P. J. (2013). Heart rate responses provide an objective evaluation of human disturbance stimuli in breeding birds. Conservation Physiology, 1(1), cot013–cot013. https://doi.org/10.1093/conphys/cot013

Ellenberg, U., Mattern, T., Seddon, P. J., & Jorquera, G. L. (2006). Physiological and reproductive consequences of human disturbance in Humboldt penguins: The need for species-specific visitor management. Biological Conservation, 133(1), 95–106. https://doi.org/10.1016/j.biocon.2006.05.019

Ellenberg, U., Setiawan, A. N., Cree, A., Houston, D. M., & Seddon, P. J. (2007). Elevated hormonal stress response and reduced reproductive output in Yellow-eyed penguins exposed to unregulated tourism. General and Comparative Endocrinology, 152(1), 54–63. https://doi.org/10.1016/j.ygcen.2007.02.022

Everard, M. (2020). Managing socio-ecological systems: who, what and how much? The case of the Banas river, Rajasthan, India. Current Opinion in Environmental Sustainability, 44, 16–25. https://doi.org/10.1016/j.cosust.2020.03.004

Fieberg, J., & Kochanny, C. O. (2005). Quantifying Home-Range Overlap: The Importance of the Utilization Distribution. The Journal of Wildlife Management, 69(4), 1346–1359. https://doi.org/10.2193/0022-541X(2005)69[1346:QHOTIO]2.0.CO;2

French, R. K., Muller, C. G., Chilvers, B. L., & Battley, P. F. (2019). Behavioural consequences of human disturbance on subantarctic Yellow-eyed Penguins Megadyptes antipodes. Bird Conservation International, 29(2), 277–290. https://doi.org/10.1017/S0959270918000096

Furness, R. W., & Camphuysen, K. (C. J.). (1997). Seabirds as monitors of the marine environment. ICES Journal of Marine Science, 54(4), 726–737. https://doi.org/10.1006/jmsc.1997.0243

Gordo, O., Brotons, L., Herrando, S., & Gargallo, G. (2021). Rapid behavioural response of urban birds to COVID-19 lockdown. Proceedings of the Royal Society B: Biological Sciences, 288(1946), 20202513. https://doi.org/10.1098/rspb.2020.2513

Grémillet, D., Pichegru, L., Kuntz, G., Woakes, A. G., Wilkinson, S., Crawford, R. J. M., & Ryan, P. G. (2008). A junk-food hypothesis for gannets feeding on fishery waste. Proceedings of the Royal Society B: Biological Sciences, 275(1639), 1149–1156. https://doi.org/10.1098/rspb.2007.1763

Hazen, E. L., Abrahms, B., Brodie, S., Carroll, G., Jacox, M. G., Savoca, M. S., Scales, K. L., Sydeman, W. J., & Bograd, S. J. (2019). Marine top predators as climate and ecosystem sentinels. Frontiers in Ecology and the Environment, 17(10), 565–574. https://doi.org/10.1002/fee.2125

Hentati-Sundberg, J., Berglund, P.-A., Hejdström, A., & Olsson, O. (2021). COVID-19 lockdown reveals tourists as seabird guardians. Biological Conservation, 254, 108950. https://doi.org/10.1016/j.biocon.2021.108950

Hijmans, R. J. (2022). raster: Geographic Data Analysis and Modeling. https://rspatial.org/raster

Hobson, K. A., Piatt, J. F., & Pitocchelli, J. (1994). Using Stable Isotopes to Determine Seabird Trophic Relationships. Journal of Animal Ecology, 63(4), 786–798. https://doi.org/10.2307/5256

Holles, S., Simpson, S. D., Radford, A. N., Berten, L., & Lecchini, D. (2013). Boat noise disrupts orientation behaviour in a coral reef fish. Marine Ecology Progress Series, 485, 295–300. https://doi.org/10.3354/meps10346

Ivanova, S., Kessel, S. T., Espinoza, M., McLean, M. F., O’Neill, C., Landry, J., Hussey, N. E., Williams, R., Vagle, S., & Fisk, A. T. (2020). Shipping alters the movement and behavior of Arctic cod (Boreogadus saida), a keystone fish in Arctic marine ecosystems. Ecological Applications. https://doi.org/10.1002/eap.2050

Jackson, A. L., Inger, R., Parnell, A. C., & Bearhop, S. (2011). Comparing isotopic niche widths among and within communities: SIBER – Stable Isotope Bayesian Ellipses in R. Journal of Animal Ecology, 80(3), 595–602. https://doi.org/10.1111/j.1365-2656.2011.01806.x

Jarrett, D., Calladine, J., Cook, A. S. C. P., Upton, A., Williams, J., Williams, S., Wilson, J. M., Wilson, M. W., Woodward, I., & Humphreys, E. M. (2021). Behavioural responses of non-breeding waterbirds to marine traffic in the near-shore environment. Bird Study, 68(4), 443–454. https://doi.org/10.1080/00063657.2022.2113855

Joly, N., Chiaradia, A., Georges, J., & Saraux, C. (2022). Environmental effects on foraging performance in little penguins: a matter of phenology and short-term variability. Marine Ecology Progress Series, 692, 151–168. https://doi.org/10.3354/meps14058

Kadykalo, A. N., Beaudoin, C., Hackenburg, D. M., Young, N., & Cooke, S. J. (2022). Social– ecological systems approaches are essential for understanding and responding to the complex impacts of COVID-19 on people and the environment. PLOS Sustainability and Transformation, 1(4), e0000006. https://doi.org/10.1371/journal.pstr.0000006

Kato, A., Ropert-Coudert, Y., & Chiaradia, A. (2008). Regulation of Trip Duration by an Inshore Forager, the Little Penguin (Eudyptula Minor), During Incubation. The Auk, 125(3), 588–593. https://doi.org/10.1525/auk.2008.06273

Klomp, N. I., & Wooller, R. D. (1991). Patterns of Arrival and Departure by Breeding Little Penguins at Penguin Island, Western Australia. Emu, 91(1), 32–35. https://doi.org/10.1071/mu9910032

Lamers, M., & Student, J. (2021). Learning from COVID-19? An environmental mobilities and flows perspective on dynamic vulnerabilities in coastal tourism settings. Maritime Studies. https://doi.org/10.1007/s40152-021-00242-1

Mace, G. M. (2014). Whose conservation? Science, 345(6204), 1558–1560. https://doi.org/10.1126/science.1254704

Manenti, R., Mori, E., Di Canio, V., Mercurio, S., Picone, M., Caffi, M., Brambilla, M., Ficetola, G. F., & Rubolini, D. (2020). The good, the bad and the ugly of COVID-19 lockdown effects on wildlife conservation: Insights from the first European locked down country. Biological Conservation, 249, 108728. https://doi.org/10.1016/j.biocon.2020.108728

Margalida, A., Colomer, M. À., & Oro, D. (2014). Man-induced activities modify demographic parameters in a long-lived species: effects of poisoning and health policies. Ecological Applications, 24(3), 436–444. https://doi.org/10.1890/13-0414.1

Mattern, T., Ellenberg, U., Houston, D. M., Lamare, M., Davis, L. S., Heezik, Y. van, & Seddon, P. J. (2013). Straight Line Foraging in Yellow-Eyed Penguins: New Insights into Cascading Fisheries Effects and Orientation Capabilities of Marine Predators. PLOS ONE, 8(12), e84381. https://doi.org/10.1371/journal.pone.0084381

McClintock, B. T., & Michelot, T. (2018). momentuHMM: R package for generalized hidden Markov models of animal movement. Methods in Ecology and Evolution, 9(6), 1518–1530. https://doi.org/10.1111/2041-210X.12995

Pelletier, L., Chiaradia, A., Kato, A., & Ropert□Coudert, Y. (2014). Fine-scale spatial age segregation in the limited foraging area of an inshore seabird species, the little penguin. Oecologia. https://doi.org/10.1007/s00442-014-3018-3

Piatt, J. F., Harding, A. M. A., Shultz, M. T., Speckman, S. G., van Pelt, T. I., Drew, G. S., & Kettle, A. B. (2007). Seabirds as indicators of marine food supplies: Cairns revisited. Marine Ecology Progress Series, 352, 221–234. USGS Publications Warehouse. https://doi.org/10.3354/meps07078

Piatt, J. F., Sydeman, W. J., & Wiese, F. (2007). Introduction: a modern role for seabirds as indicators. Marine Ecology Progress Series, 352, 199–204. https://www.int-res.com/abstracts/meps/v352/p199-204/

Pichegru, L., Grémillet, D., Crawford, R. J. M., & Ryan, P. G. (2010). Marine no-take zone rapidly benefits endangered penguin. Biology Letters. https://doi.org/10.1098/rsbl.2009.0913

Pichegru, L., Nyengera, R., McInnes, A. M., & Pistorius, P. (2017). Avoidance of seismic survey activities by penguins. Scientific Reports, 7(1), 16305. https://doi.org/10.1038/s41598-017-16569-x

Pichegru, L., Vibert, L., Thiebault, A., Charrier, I., Stander, N., Ludynia, K., Lewis, M., Carpenter-Kling, T., & McInnes, A. (2022). Maritime traffic trends around the southern tip of Africa – Did marine noise pollution contribute to the local penguins’ collapse? Science of The Total Environment, 849, 157878. https://doi.org/10.1016/j.scitotenv.2022.157878

Pinheiro, J., Bates, D., DebRoy, S., Sarkar, D., & R Core Team. (2021). nlme: Linear and Nonlinear Mixed Effects Models. https://CRAN.R-project.org/package=nlme

Post, D. M., Layman, C. A., Arrington, D. A., Takimoto, G., Quattrochi, J., & Montaña, C. G. (2007). Getting to the fat of the matter: models, methods and assumptions for dealing with lipids in stable isotope analyses. Oecologia, 152(1), 179–189. https://doi.org/10.1007/s00442-006-0630-x

Poupart, T. A., Waugh, S. M., Bost, C., Bost, C.-A., Dennis, T., Lane, R., Rogers, K., Sugishita, J., Taylor, G. A., Wilson, K.-J., Zhang, J., & Arnould, J. P. Y. (2017). Variability in the foraging range of Eudyptula minor across breeding sites in central New Zealand. New Zealand Journal of Zoology, 44(3), 225–244. https://doi.org/10.1080/03014223.2017.1302970

R Core Team. (2023). R: A Language and Environment for Statistical Computing. R Foundation for Statistical Computing. https://www.R-project.org/

Ramírez, F., Afán, I., Davis, L. S., & Chiaradia, A. (2017). Climate impacts on global hot spots of marine biodiversity. Science Advances, 3(2), e1601198. https://doi.org/10.1126/sciadv.1601198

Ramírez, F., Forero, M. G., Hobson, K. A., & Chiaradia, A. (2015). Older female little penguins Eudyptula minor adjust nutrient allocations to both eggs. Journal of Experimental Marine Biology and Ecology, 468, 91–96. https://doi.org/10.1016/j.jembe.2015.03.020

Ratcliffe, N., Bell, M., Pelembe, T., Boyle, D., Benjamin, R., White, R., Godley, B., Stevenson, J., & Sanders, S. (2010). The eradication of feral cats from Ascension Island and its subsequent recolonization by seabirds. Oryx, 44(01), 20–29. https://doi.org/10.1017/S003060530999069X

Rodríguez, A., Chiaradia, A., Wasiak, P., Renwick, L., & Dann, P. (2016). Waddling on the Dark Side: Ambient Light Affects Attendance Behavior of Little Penguins. Journal of Biological Rhythms, 31(2), 194–204. https://doi.org/10.1177/0748730415626010

Rodríguez, A., Holmberg, R., Dann, P., & Chiaradia, A. (2018). Penguin colony attendance under artificial lights for ecotourism. Journal of Experimental Zoology Part A: Ecological and Integrative Physiology, 329(8–9), 457–464. https://doi.org/10.1002/jez.2155

Rutz, C., Loretto, M.-C., Bates, A. E., Davidson, S. C., Duarte, C. M., Jetz, W., Johnson, M., Kato, A., Kays, R., Mueller, T., Primack, R. B., Ropert-Coudert, Y., Tucker, M. A., Wikelski, M., & Cagnacci, F. (2020). COVID-19 lockdown allows researchers to quantify the effects of human activity on wildlife. Nature Ecology & Evolution, 4(9), 1156–1159. https://doi.org/10.1038/s41559-020-1237-z

Sala, M. M., Peters, F., Sebastián, M., Cardelús, C., Calvo, E., Marrasé, C., Massana, R., Pelejero, C., Sala-Coromina, J., Vaqué, D., & Gasol, J. M. (2021). COVID-19 lockdown moderately increased oligotrophy at a marine coastal site. Science of The Total Environment, 151443. https://doi.org/10.1016/j.scitotenv.2021.151443

Salton, M., Saraux, C., Dann, P., & Chiaradia, A. (2015). Carry-over body mass effect from winter to breeding in a resident seabird, the little penguin. Royal Society Open Science. https://doi.org/10.1098/rsos.140390

Sánchez, S., Reina, R. D., Kato, A., Ropert-Coudert, Y., Cavallo, C., Hays, G. C., & Chiaradia, A. (2018). Within-colony spatial segregation leads to foraging behaviour variation in a seabird. Marine Ecology Progress Series, 606, 215–230. https://doi.org/10.3354/meps12764

Saraux, C., Chiaradia, A., Maho, Y., & RopertDCoudert, Y. (2011). Everybody needs somebody: unequal parental effort in little penguins. https://doi.org/10.1093/BEHECO/ARR049

Seaman, D. E., Griffith, B., & Powell, R. A. (1998). KERNELHR: A program for estimating animal home ranges. Wildlife Society Bulletin, 26(1), 95–100. USGS Publications Warehouse. http://pubs.er.usgs.gov/publication/1015784

Siddig, A. A. H., Ellison, A. M., Ochs, A., Villar-Leeman, C., & Lau, M. K. (2016). How do ecologists select and use indicator species to monitor ecological change? Insights from 14 years of publication in Ecological Indicators. Ecological Indicators, 60, 223–230. https://doi.org/10.1016/j.ecolind.2015.06.036

Sugihara, G., May, R., Ye, H., Hsieh, C., Deyle, E., Fogarty, M., & Munch, S. (2012). Detecting Causality in Complex Ecosystems. Science, 338(6106), 496–500. https://doi.org/10.1126/science.1227079

Sutherland, D. R., & Dann, P. (2014). Population trends in a substantial colony of Little Penguins: three independent measures over three decades. Biodiversity and Conservation, 23(1), 241–250. https://doi.org/10.1007/s10531-013-0597-y

Trathan, P. N., GarcíaDBorboroglu, P., Boersma, D., Bost, C., Crawford, R. J. M., Crossin, G. T., Cuthbert, R. J., Dann, P., Davis, L. S., De La Puente, S., Ellenberg, U., Lynch, H. J., Mattern, T., Pütz, K., Seddon, P. J., Trivelpiece, W., & Wienecke, B. (2015). Pollution, habitat loss, fishing, and climate change as critical threats to penguins. Conservation Biology, 29(1), 31–41. https://doi.org/10.1111/cobi.12349

Tucker, M. A., Alexandrou, O., Bierregaard Jr., R. O., Bildstein, K. L., Böhning-Gaese, K., Bracis, C., Brzorad, J. N., Buechley, E. R., Cabot, D., Calabrese, J. M., Carrapato, C., Chiaradia, A., Davenport, L. C., Davidson, S. C., Desholm, M., DeSorbo, C. R., Domenech, R., Enggist, P., Fagan, W. F., … Mueller, T. (2019). Large birds travel farther in homogeneous environments. Global Ecology and Biogeography, 28(5), 576–587. https://doi.org/10.1111/geb.12875

Tucker, M. A., Böhning-Gaese, K., Fagan, W. F., Fryxell, J. M., Van Moorter, B., Alberts, S. C., Ali, A. H., Allen, A. M., Attias, N., Avgar, T., Bartlam-Brooks, H., Bayarbaatar, B., Belant, J. L., Bertassoni, A., Beyer, D., Bidner, L., van Beest, F. M., Blake, S., Blaum, N., … Mueller, T. (2018). Moving in the Anthropocene: Global reductions in terrestrial mammalian movements. Science, 359(6374), 466–469. https://doi.org/10.1126/science.aam9712

Viblanc, V. A., Smith, A. D., Gineste, B., & Groscolas, R. (2012). Coping with continuous human disturbance in the wild: insights from penguin heart rate response to various stressors. BMC Ecology, 12(1), 10. https://doi.org/10.1186/1472-6785-12-10

Watanuki, Y., Wanless, S., Harris, M., Lovvorn, J. R., Miyazaki, M., Tanaka, H., & Sato, K. (2006). Swim speeds and stroke patterns in wing-propelled divers: a comparison among alcids and a penguin. Journal of Experimental Biology, 209(7), 1217–1230. https://doi.org/10.1242/jeb.02128

Wei, Y., Wu, S., & Tesemma, Z. (2018). Re-orienting technological development for a more sustainable human–environmental relationship. Current Opinion in Environmental Sustainability, 33, 151–160. https://doi.org/10.1016/j.cosust.2018.05.022

Wilson, R. P., Pütz, K., Peters, G., Culik, B., Scolaro, J. A., Charrassin, J.-B., & Ropert-Coudert, Y. (1997). Long-Term Attachment of Transmitting and Recording Devices to Penguins and Other Seabirds. Wildlife Society Bulletin (1973–2006), 25(1), 101–106. https://www.jstor.org/stable/3783290

