## Supplementary tables and figure for "COVID-related anthropause highlights the impact of marine traffic on breeding little penguins"

**Supplementary Information**

Supplementary table 1. Model selection for (A) Daily number of spectators (n = 1342), (B) Daily number of vessels (n = 824), (C) Overlap among penguins (n = 110), (D) Overlap between penguins and marine traffic (n = 49), (E) Departure time relative to nautical dawn (n = 21), (F) Arrival time relative to nautical dusk (n = 21), (G) Mass gain per day at sea (n = 21), (H) Chick meal size (n = 14).

(A)

| month | year | *df* | logLik | AICc | delta | weight |
| --- | --- | --- | --- | --- | --- | --- |
| + | + | 15 | -10,411.41 | 20,853.18 | 0.00 | 1.00 |
|  | + | 12 | -10,599.91 | 21,224.05 | 370.87 | 0.00 |
| + |  | 5 | -10,809.75 | 21,629.54 | 776.36 | 0.00 |
|  |  | 2 | -10,920.32 | 21,844.65 | 991.46 | 0.00 |

(B)

| month | year | *df* | logLik | AICc | delta | weight |
| --- | --- | --- | --- | --- | --- | --- |
| + | + | 11 | -4,677.40 | 9,377.13 | 0.00 | 1.00 |
|  | + | 8 | -4,706.24 | 9,428.66 | 51.53 | 0.00 |
| + |  | 5 | -4,722.13 | 9,454.33 | 77.20 | 0.00 |
|  |  | 2 | -4,747.70 | 9,499.41 | 122.28 | 0.00 |

(C)

| Year_A | Year_B | *df* | logLik | AICc | delta | weight |
| --- | --- | --- | --- | --- | --- | --- |
| + | + | 22 | 24.25 | 7.14 | 0.00 | 1.00 |
| + |  | 12 | -0.64 | 28.49 | 21.35 | 0.00 |
|  | + | 12 | -0.64 | 28.49 | 21.35 | 0.00 |
|  |  | 2 | -15.50 | 35.12 | 27.98 | 0.00 |

(D)

| boat_year | penguin_year | *df* | logLik | AICc | delta | weight |
| --- | --- | --- | --- | --- | --- | --- |
|  | + | 8 | 89.82 | -160.05 | 0.00 | 1.00 |
| + | + | 14 | 93.95 | -147.55 | 12.50 | 0.00 |
|  |  | 2 | 20.95 | -37.63 | 122.42 | 0.00 |
| + |  | 8 | 21.39 | -23.17 | 136.88 | 0.00 |

(E)

| breeding_stage | mean_boat | mean_spec | season | *df* | logLik | AICc | delta | weight |
| --- | --- | --- | --- | --- | --- | --- | --- | --- |
| + |  |  |  | 4 | -74.65 | 159.81 | 0.00 | 0.70 |
| + |  | -1.50 |  | 5 | -74.47 | 162.94 | 3.13 | 0.15 |
| + | 1.17 |  |  | 5 | -74.62 | 163.23 | 3.42 | 0.13 |
| + | 2.25 | -1.93 |  | 6 | -74.34 | 166.68 | 6.87 | 0.02 |
| + |  |  | + | 10 | -68.09 | 178.18 | 18.37 | 0.00 |
| + |  | -7.94 | + | 11 | -67.65 | 186.63 | 26.82 | 0.00 |
| + | -4.54 |  | + | 11 | -67.95 | 187.24 | 27.43 | 0.00 |
|  |  |  |  | 2 | -92.46 | 189.59 | 29.78 | 0.00 |
|  | -2.94 |  |  | 3 | -92.42 | 192.24 | 32.43 | 0.00 |
|  |  | -1.55 |  | 3 | -92.43 | 192.26 | 32.45 | 0.00 |
|  | -2.30 | -1.09 |  | 4 | -92.40 | 195.30 | 35.49 | 0.00 |
| + | 0.04 | -7.96 | + | 12 | -67.65 | 198.30 | 38.49 | 0.00 |
|  |  |  | + | 8 | -91.53 | 211.05 | 51.24 | 0.00 |
|  | -29.13 |  | + | 9 | -90.71 | 215.79 | 55.98 | 0.00 |
|  |  | -5.92 | + | 9 | -91.48 | 217.33 | 57.52 | 0.00 |
|  | -42.81 | 17.81 | + | 10 | -90.46 | 222.91 | 63.10 | 0.00 |

(F)

| breeding_stage | mean_boat | mean_spec | season | *df* | logLik | AICc | delta | weight |
| --- | --- | --- | --- | --- | --- | --- | --- | --- |
|  |  |  |  | 2 | -78.57 | 161.81 | 0.00 | 0.53 |
|  |  | -1.34 |  | 3 | -78.46 | 164.34 | 2.53 | 0.15 |
|  | 1.64 |  |  | 3 | -78.52 | 164.45 | 2.64 | 0.14 |
|  |  |  | + | 8 | -69.12 | 166.25 | 4.44 | 0.06 |
| + |  |  |  | 4 | -77.96 | 166.42 | 4.61 | 0.05 |
|  | 2.74 | -1.88 |  | 4 | -78.33 | 167.16 | 5.35 | 0.04 |
| + |  | -1.68 |  | 5 | -77.79 | 169.57 | 7.76 | 0.01 |
| + | 1.57 |  |  | 5 | -77.91 | 169.82 | 8.01 | 0.01 |
|  | 7.59 |  | + | 9 | -68.75 | 171.86 | 10.05 | 0.00 |
|  |  | -3.00 | + | 9 | -69.02 | 172.40 | 10.59 | 0.00 |
| + | 2.78 | -2.19 |  | 6 | -77.64 | 173.28 | 11.47 | 0.00 |
| + |  |  | + | 10 | -67.55 | 177.10 | 15.29 | 0.00 |
|  | 15.97 | -10.23 | + | 10 | -67.92 | 177.84 | 16.04 | 0.00 |
| + |  | -11.66 | + | 11 | -66.47 | 184.28 | 22.47 | 0.00 |
| + | 8.72 |  | + | 11 | -67.10 | 185.54 | 23.73 | 0.00 |
| + | 19.56 | -19.85 | + | 12 | -64.39 | 191.78 | 29.97 | 0.00 |

(G)

| breeding_stage | mean_boat | mean_spec | season | *df* | logLik | AICc | delta | weight |
| --- | --- | --- | --- | --- | --- | --- | --- | --- |
| + | -56.86 |  |  | 5 | -101.14 | 216.28 | 0.00 | 0.74 |
| + | -50.32 | -10.99 |  | 6 | -100.43 | 218.87 | 2.59 | 0.20 |
| + |  | -20.64 |  | 5 | -104.48 | 222.97 | 6.69 | 0.03 |
| + |  |  |  | 4 | -106.30 | 223.09 | 6.81 | 0.02 |
| + | -106.91 |  | + | 11 | -88.48 | 228.30 | 12.02 | 0.00 |
| + |  |  | + | 10 | -95.90 | 233.80 | 17.52 | 0.00 |
| + |  | -58.24 | + | 11 | -94.28 | 239.90 | 23.62 | 0.00 |
| + | -106.20 | -1.39 | + | 12 | -88.48 | 239.96 | 23.69 | 0.00 |
|  |  |  |  | 2 | -121.76 | 248.18 | 31.90 | 0.00 |
|  | -26.95 |  |  | 3 | -121.53 | 250.48 | 34.20 | 0.00 |
|  |  | -13.70 |  | 3 | -121.58 | 250.57 | 34.30 | 0.00 |
|  | -21.09 | -9.38 |  | 4 | -121.46 | 253.42 | 37.14 | 0.00 |
|  |  |  | + | 8 | -120.12 | 268.24 | 51.96 | 0.00 |
|  | 74.71 |  | + | 9 | -119.76 | 273.89 | 57.61 | 0.00 |
|  |  | 55.29 | + | 9 | -119.88 | 274.13 | 57.85 | 0.00 |
|  | 60.20 | 20.65 | + | 10 | -119.74 | 281.49 | 65.21 | 0.00 |

(H)

| breeding_stage | mean_boat | mean_spec | season | *df* | logLik | AICc | delta | weight |
| --- | --- | --- | --- | --- | --- | --- | --- | --- |
|  | -22.78 |  |  | 3 | -60.54 | 129.47 | 0.00 | 0.32 |
|  |  | -12.88 |  | 3 | -60.85 | 130.09 | 0.62 | 0.24 |
|  |  |  |  | 2 | -62.61 | 130.32 | 0.85 | 0.21 |
|  | -16.82 | -8.59 |  | 4 | -59.72 | 131.89 | 2.41 | 0.10 |
| + | -23.41 |  |  | 4 | -60.44 | 133.33 | 3.85 | 0.05 |
| + |  |  |  | 3 | -62.61 | 133.62 | 4.14 | 0.04 |
| + |  | -13.19 |  | 4 | -60.77 | 133.99 | 4.52 | 0.03 |
| + | -17.41 | -8.86 |  | 5 | -59.56 | 136.62 | 7.15 | 0.01 |
|  |  |  | + | 8 | -50.60 | 146.00 | 16.52 | 0.00 |
|  | -9.62 |  | + | 9 | -50.50 | 164.00 | 34.53 | 0.00 |
| + |  |  | + | 9 | -50.57 | 164.13 | 34.66 | 0.00 |
|  |  | -2.80 | + | 9 | -50.58 | 164.15 | 34.68 | 0.00 |
| + | -20.34 |  | + | 10 | -50.29 | 193.91 | 64.44 | 0.00 |
| + |  | -6.60 | + | 10 | -50.48 | 194.30 | 64.82 | 0.00 |
|  | -10.57 | 0.98 | + | 10 | -50.50 | 194.33 | 64.86 | 0.00 |
| + | -18.74 | -2.41 | + | 11 | -50.28 | 254.56 | 125.08 | 0.00 |

Supplementary table 2. Best model summary for (A) Daily number of spectators (n = 1342), (B) Daily number of vessels (n = 824), (C) Overlap among penguins (n = 110), (D) Overlap between penguins and marine traffic (n = 49), (E) Departure time relative to nautical dawn (n = 21), (F) Arrival time relative to nautical dusk (n = 21), (G) Mass gain per day at sea (n = 21), (H) Chick meal size (n = 14).

(A)

| Parameter | *b* | *t* | *df* | *p* | β | 95% CI (β) |
| --- | --- | --- | --- | --- | --- | --- |
| (Intercept) | 1,134.43 | 19.46 | 1,328 | < .001 | -0.59 | [-0.73, -0.46] |
| month [10] | 88.50 | 2.01 | 1,328 | .044 | 0.11 | [0.00, 0.21] |
| month [11] | 232.16 | 5.24 | 1,328 | < .001 | 0.28 | [0.18, 0.39] |
| month [12] | 823.10 | 18.72 | 1,328 | < .001 | 0.99 | [0.89, 1.10] |
| year [2011] | 18.64 | 0.26 | 1,328 | .798 | 0.02 | [-0.15, 0.20] |
| year [2012] | 114.07 | 1.56 | 1,328 | .118 | 0.14 | [-0.03, 0.31] |
| year [2013] | 215.03 | 2.95 | 1,328 | .003 | 0.26 | [0.09, 0.43] |
| year [2014] | 292.11 | 4.01 | 1,328 | < .001 | 0.35 | [0.18, 0.53] |
| year [2015] | 552.59 | 7.58 | 1,328 | < .001 | 0.67 | [0.49, 0.84] |
| year [2016] | 656.29 | 9.00 | 1,328 | < .001 | 0.79 | [0.62, 0.97] |
| year [2017] | 556.21 | 7.63 | 1,328 | < .001 | 0.67 | [0.50, 0.84] |
| year [2018] | 601.84 | 8.26 | 1,328 | < .001 | 0.73 | [0.55, 0.90] |
| year [2019] | 461.52 | 6.33 | 1,328 | < .001 | 0.56 | [0.38, 0.73] |
| year [2020] | -1,242.79 | -17.05 | 1,328 | < .001 | -1.50 | [-1.67, -1.33] |

(B)

| Parameter | *b* | *t* | *df* | *p* | β | 95% CI (β) |
| --- | --- | --- | --- | --- | --- | --- |
| (Intercept) | 206.42 | 26.85 | 814 | < .001 | -0.70 | [-0.90, -0.50] |
| month [11] | 15.69 | 2.16 | 814 | .031 | 0.20 | [0.02, 0.39] |
| month [12] | 27.99 | 4.10 | 814 | < .001 | 0.36 | [0.19, 0.54] |
| month [9] | -22.46 | -3.27 | 814 | .001 | -0.29 | [-0.47, -0.12] |
| year [2015] | 42.90 | 4.71 | 814 | < .001 | 0.56 | [0.33, 0.79] |
| year [2016] | 49.94 | 5.49 | 814 | < .001 | 0.65 | [0.42, 0.88] |
| year [2017] | 57.35 | 6.30 | 814 | < .001 | 0.75 | [0.51, 0.98] |
| year [2018] | 84.89 | 9.33 | 814 | < .001 | 1.10 | [0.87, 1.33] |
| year [2019] | 58.50 | 5.89 | 814 | < .001 | 0.76 | [0.51, 1.01] |
| year [2020] | 50.61 | 5.56 | 814 | < .001 | 0.66 | [0.43, 0.89] |

(C)

| Parameter | *b* | *t* | *df* | *p* | β | 95% CI (β) |
| --- | --- | --- | --- | --- | --- | --- |
| (Intercept) | 0.51 | 3.86 | 29 | < .001 | 0.51 | [0.26, 0.77] |
| year A [2015] | 1.23 | 5.98 | 29 | < .001 | 1.23 | [0.84, 1.65] |
| year A [2016] | 0.24 | 1.79 | 29 | .074 | 0.24 | [-0.02, 0.51] |
| year A [2017] | 0.32 | 2.30 | 29 | .021 | 0.32 | [0.05, 0.60] |
| year A [2018] | 0.68 | 4.18 | 29 | < .001 | 0.68 | [0.37, 1.01] |
| year A [2019] | 0.42 | 2.90 | 29 | .004 | 0.42 | [0.14, 0.71] |
| year A [2020] | 0.23 | 1.67 | 29 | .095 | 0.23 | [-0.04, 0.49] |
| year B [2015] | 1.23 | 5.98 | 29 | < .001 | 1.23 | [0.84, 1.65] |
| year B [2016] | 0.24 | 1.79 | 29 | .074 | 0.24 | [-0.02, 0.51] |
| year B [2017] | 0.32 | 2.30 | 29 | .021 | 0.32 | [0.05, 0.60] |
| year B [2018] | 0.68 | 4.18 | 29 | < .001 | 0.68 | [0.37, 1.01] |
| year B [2019] | 0.42 | 2.90 | 29 | .004 | 0.42 | [0.14, 0.71] |
| year B [2020] | 0.23 | 1.67 | 29 | .095 | 0.23 | [-0.04, 0.49] |

(D)

| Parameter | *b* | *t* | *df* | *p* | β | 95% CI (β) |
| --- | --- | --- | --- | --- | --- | --- |
| (Intercept) | 6.68 | 12.45 | 42 | < .001 | 6.68 | [5.68, 7.78] |
| penguin year [2015] | 40.61 | 10.59 | 42 | < .001 | 40.61 | [33.47, 48.52] |
| penguin year [2016] | -0.42 | -0.58 | 42 | .563 | -0.42 | [-1.87, 1.01] |
| penguin year [2017] | -3.88 | -6.68 | 42 | < .001 | -3.88 | [-5.07, -2.79] |
| penguin year [2018] | -5.14 | -9.33 | 42 | < .001 | -5.14 | [-6.27, -4.11] |
| penguin year [2019] | -2.00 | -3.06 | 42 | .002 | -2.00 | [-3.31, -0.74] |
| penguin year [2020] | -1.26 | -1.82 | 42 | .069 | -1.26 | [-2.63, 0.09] |

(E)

| Parameter | *b* | *t* | *df* | *p* | β | 95% CI (β) |
| --- | --- | --- | --- | --- | --- | --- |
| (Intercept) | -74.94 | -21.68 | 18 | < .001 | -1.17 | [-1.53, -0.81] |
| breeding stage [Incubation] | 43.16 | 8.83 | 18 | < .001 | 2.13 | [1.62, 2.64] |
| breeding stage [PG] | 27.80 | 5.69 | 18 | < .001 | 1.37 | [0.87, 1.88] |

(F)

| Parameter | *b* | *t* | *df* | *p* | β | 95% CI (β) |
| --- | --- | --- | --- | --- | --- | --- |
| (Intercept) | 9.25 | 4.06 | 20 | .001 | -0.00 | [-0.46, 0.46] |

(G)

| Parameter | *b* | *t* | *df* | *p* | β | 95% CI (β) |
| --- | --- | --- | --- | --- | --- | --- |
| (Intercept) | 302.43 | 24.05 | 17 | < .001 | 0.87 | [0.54, 1.19] |
| breeding stage [Incubation] | -173.21 | -9.68 | 17 | < .001 | -2.12 | [-2.58, -1.66] |
| breeding stage [PG] | -39.24 | -2.20 | 17 | .042 | -0.48 | [-0.94, -0.02] |
| mean boat | -56.86 | -3.28 | 17 | .004 | -0.31 | [-0.50, -0.11] |

(H)

| Parameter | *b* | *t* | *df* | *p* | β | 95% CI (β) |
| --- | --- | --- | --- | --- | --- | --- |
| (Intercept) | 273.45 | 51.12 | 12 | < .001 | -0.00 | [-0.52, 0.52] |
| mean boat | -22.78 | -2.04 | 12 | .064 | -0.51 | [-1.05, 0.04] |

Supplementary table 3. Stable isotopes samples distribution from 2010 to 2020

| Breeding stage | sex | 2010 | 2011 | 2012 | 2013 | 2014 | 2015 | 2016 | 2017 | 2018 | 2019 | 2020 |
| --- | --- | --- | --- | --- | --- | --- | --- | --- | --- | --- | --- | --- |
| Guard |  | 22 | 20 | 13 | 13 | 20 | 14 | 19 | 20 | 19 | 20 | 17 |
|  | F | 9 | 9 | 1 | 3 | 7 | 8 | 9 | 11 | 10 | 10 | 8 |
|  | M | 6 | 9 | 8 |  | 6 | 7 | 10 | 10 | 10 | 10 | 9 |
| Incubation | F | 9 | 10 |  | 7 | 9 | 11 | 12 | 11 | 9 | 10 | 9 |
|  | M | 11 | 10 |  | 9 | 10 | 9 | 8 | 10 | 10 | 11 | 11 |
| Post-guard |  | 20 | 20 | 20 |  | 14 | 19 | 20 | 23 | 9 | 19 | 18 |
|  | F | 9 | 6 | 2 |  |  |  |  | 9 | 4 | 9 | 9 |
|  | M | 8 | 10 | 2 |  |  |  |  | 6 | 5 | 9 | 9 |

Supplementary figure 1. Evolution of the proportion of vessel types in the Bass strait marine traffic. Vessels types representing less than 0.5% of the dataset were merged in the “Other” type.

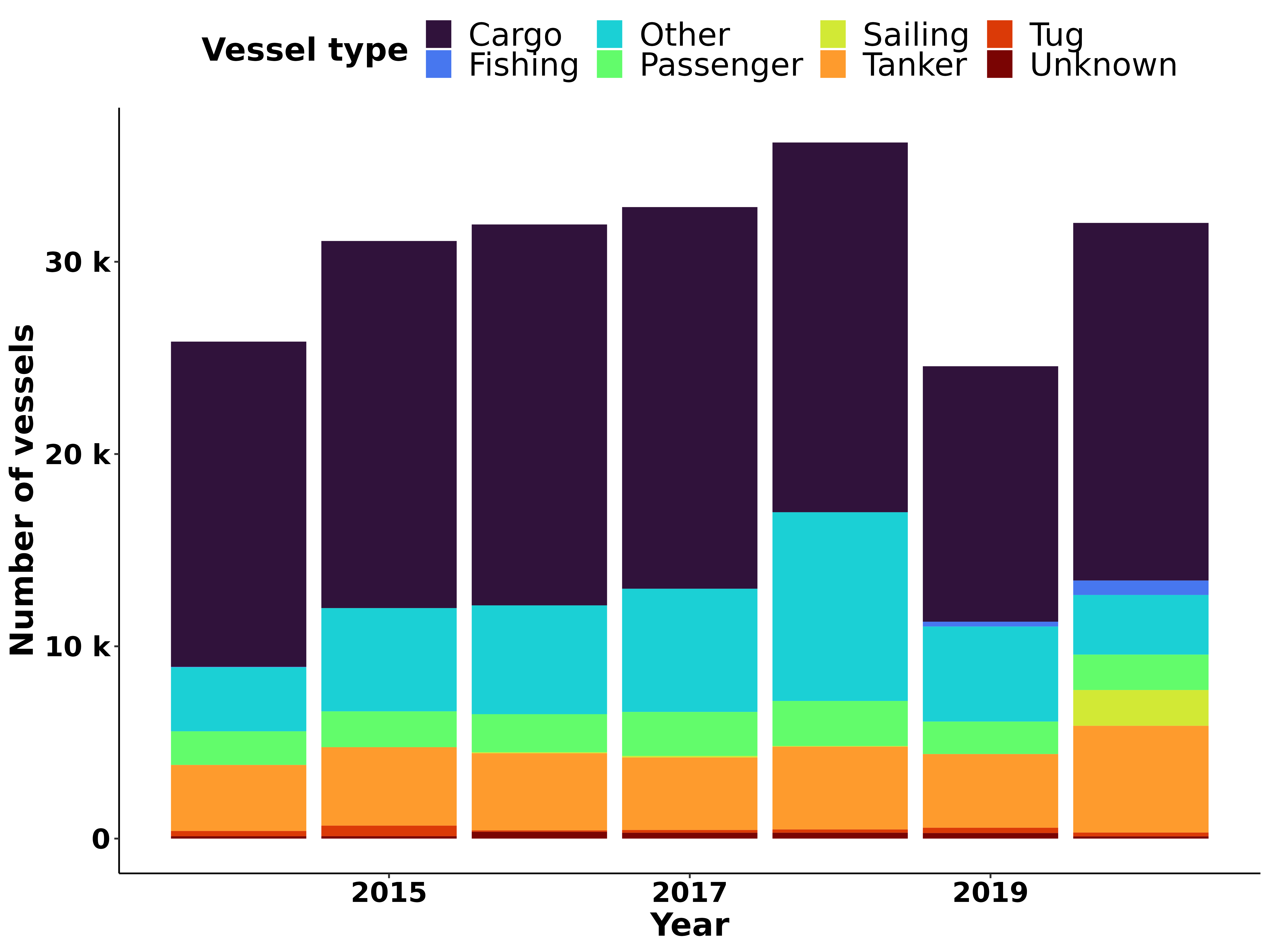
